# Model evaluation metrics of Virtual Cell Challenge 2026

**DOI:** 10.64898/2026.11.17.767214

**Authors:** Alexander Dobin, Abhinav Adduri, Jeremy Sullivan, Jonathan Lyda, Reshma Mehta, Dara E. Leto, Paul Datlinger, Sadie VanHorn, Shahab Chizari, Ninad Ranadive, Sanmati Nagesh, Po-Yuan Tung, Nianzhen Li, Joseph Caputo, Brian S. Plosky, Chiara Ricci-Tam, Rachel Senturia, Kevin Feng, Tiffany Chu, Maggie Tsui, Basak Eraslan, Beatrice Bevilacqua, Christopher Carpenter, Rishi Verma, Zichao Yan, Joseph Min, Furkan M. Torun, Inna Averbukh, Thomas Sandmann, T.J. Chen, Severin Dicks, Ohad Mosafi, Timur Rvachov, Heidi Shin, Laksshman Sundaram, Dawn Voss, Yang Zhang, Greg Zynda, Megan van Overbeek, Usman Muzaffar, Alden Woodrow, Dave P. Burke, Patrick D. Hsu, Silvana Konermann, Arc Virtual Cell Team, Hani Goodarzi

**Affiliations:** Arc Institute, Palo Alto, CA, USA; NVIDIA Corporation, Santa Clara, CA, USA; Department of Pathology, Stanford University School of Medicine, Stanford, CA, USA; Department of Biochemistry, Stanford University School of Medicine, Stanford, CA, USA; University of California, San Francisco, San Francisco, CA, USA

## Abstract

Virtual cell models aim to predict the effects of genetic, environmental, chemical and other perturbations in silico, reducing the dependency on complex and costly experiments. The Virtual Cell Challenge 2026 motivates the field toward this goal, tasking models with zero-shot prediction of the transcriptional responses to CRISPRi knockdowns in six human cell lines. The faithfulness of a model to the experiment is assessed by metrics that compare its predictions with the real data, and designing such metrics is not trivial. We developed the metrics of the 2026 Challenge around three principles: a metric should (i) reflect the biological information extracted from experiments, (ii) be calibrated on a repeat of the experiment, and (iii) resist shortcut solutions that raise the score without predicting the response. Here we present the six resulting metrics, the reasoning behind each of them, and their evaluations on the Challenge data. Two of the metrics compare the pseudobulk expression profiles of predicted and measured cells, while four assess the genes that respond to each perturbation, the central readout of a Perturb-seq experiment. The metrics are implemented in cell-eval2, an open-source Python package. On the Challenge data, all six metrics score a repeat of the experiment above the mean baseline, and the shortcut solutions that the metrics of the 2025 Challenge left open can no longer raise the score. The new metrics are therefore more robust than the previous ones.

## 1. Introduction

Virtual cell models aim to predict how a cell responds to a perturbation where this response was never measured, especially where such measurements are not yet feasible. They become valuable to human health once they predict the response in humans at least as well as the best current preclinical models do. While cellular responses to perturbations include changes in many biological layers (for example, in the epigenome, transcriptome, proteome, and metabolome), here we focus on the changes in the transcriptome, as measured by Perturb-seq. Perturb-seq combines pooled CRISPR perturbations with single-cell RNA sequencing and measures the responses to thousands of perturbations in one experiment (Replogle et al., 2022; Nadig et al., 2025). Such data provide both the training data and the ground truth for these models. The Virtual Cell Challenge was launched to compare these models within one evaluation framework, on datasets generated for this purpose, and to accelerate their development (Roohani et al., 2025). For the 2026 Challenge, participants had to provide zero-shot predictions of the responses to CRISPR-interference (CRISPRi) knockdowns in six human cell lines (Arc Institute, 2026a; Arc Virtual Cell Initiative Team and Goodarzi, 2026).

Assessing predictive models depends heavily on the metrics that compare the predictions with experimental measurements, and designing these metrics is not trivial. Single-cell measurements are noisy, so even a repeat of the experiment does not reproduce the measured profiles exactly. A metric that ignores this noise can therefore rank a no-change prediction (i.e. the control cells) above a repeat of the experiment. In addition, a metric can respond to properties of a prediction that are unrelated to the biology of the response, such as the scale of the predicted values. In a competition, such weaknesses matter even more, since the participants optimize the score as part of their design goals. Any part of the score that can be gained without predicting the response invites what we call “shortcut solutions”. We borrow the term from the “shortcut learning” of machine learning models, which can learn decision rules that perform well on standard benchmarks but fail to transfer to more challenging conditions (Geirhos et al., 2020).

Recent benchmarks showed how strongly the conclusions about these models depend on their evaluation. Deep-learning and foundation models of single cells did not outperform deliberately simple baselines, such as the mean of the training examples or a linear model, in predicting the responses to genetic perturbations on conventional metrics (Ahlmann-Eltze et al., 2025; Csendes et al., 2025). Common metrics also overestimate performance because they are susceptible to the systematic differences between perturbed and control cells (Viñas Torné et al., 2026). Other studies introduced rank-based metrics (Wu et al., 2024) and showed how the distance and the scaling of the data change the perturbation discrimination score (Liu et al., 2025). Proposed evaluation principles include calibrating metrics against empirical reference points (Vollenweider and Bühlmann, 2026), such as a lower reference and a positive reference made by splitting the cells of the experiment (Schäfer et al., 2026).

The 2025 Challenge scored the predictions with three metrics: the perturbation discrimination with the L1 distance, the mean absolute error of the expression profiles, and the overlap of the predicted and measured differentially expressed (DE) genes (Roohani et al., 2025). However, these metrics left room for shortcut solutions (Section 2.3), and the mean absolute error scores a repeat of the experiment worse than a no-change prediction (Section 4.2). For the 2026 Challenge, we redesigned the metrics, guided by three principles: a metric should (i) reflect the biological information extracted from experiments, (ii) be calibrated on a repeat of the experiment, and (iii) resist shortcut solutions.

We present the six metrics that score the 2026 Virtual Cell Challenge, the reasoning behind each of them, and their values on the Challenge datasets. Two of them compare expression profiles (perturbation discrimination and expression error), and four compare the DE genes (fold-change error, significant-set agreement, direction fidelity yield and direction reach). We implemented these metrics in cell-eval2, an open-source Python package (Supplementary Note 1).

## 2. Data and task

### 2.1 The task

The 2026 Challenge asks for zero-shot prediction across cell lines. The responses to CRISPRi knockdowns were measured by Perturb-seq in six human cell lines from different tissues of origin and were all withheld as the ground truth (Arc Institute, 2026a; Arc Virtual Cell Initiative Team and Goodarzi, 2026). For each line, the participants received only the expression profiles of cells carrying non-targeting guides, which describe the unperturbed state of the line, and the list of genes whose knockdown they had to predict. Importantly, there was no Challenge-specific training set, and the participants could train their models on any other data, public or their own. This is the main change from the 2025 Challenge, when the prediction stayed within one cell type, and a training set measured in that same cell type was provided.

Cell lines A/B/C formed the validation phase, scored on a live leaderboard, while test lines D/E/F decided the final ranking. Each submission received a single score, the mean of the six metric scores averaged over the cell lines (Section 3.2).

### 2.2 Data and panel

The six lines were screened with the same pooled CRISPRi guide library, and profiled with 10x Genomics Flex, a probe-based single-cell assay (Arc Institute, 2026a, 2026b; Swinderman et al., 2026). All released and scored data share one axis of 18,533 genes. Each line is scored on a panel of 300 perturbations, one per target gene. The three validation lines (A/B/C) share one panel, while the three test lines (D/E/F) share another, with 129 genes in both panels. We excluded all genes targeted in any of the four essential-gene Perturb-seq screens of Replogle et al. (2022) and Nadig et al. (2025), so that no measured response to a panel perturbation could be carried over from these public datasets. Supplementary Fig. S1 shows the cumulative distributions of the target genes’ expression in the control cells and of their knockdown.

Each panel gene is represented by one dual-guide construct, which carries two guides against that gene (Replogle et al., 2020), the same in every line in which the gene is scored. The six lines differ in their native depth (median ~21,500–31,600 unique molecular identifiers (UMIs) per cell) and in the number of cells per construct (median 445–590). From the cells of each panel construct, we subsampled exactly 400 cells per line. The control pool of every line contains the same 46 non-targeting constructs, also with 400 cells each, i.e. 18,400 control cells per line. To match the sequencing depth across the lines, we thinned the counts of every cell binomially, so that each line has a median of ~20,000 UMIs per cell. We fixed the number of cells and the depth because every metric depends on both. More cells and reads add power to the Wilcoxon test that selects the genes of the DE metrics and reduce the noise in the pseudobulk profiles of the expression metrics.

### 2.3 Submissions: single cells, counts and measured controls

We require predicted single cells (cell × gene count matrix, 400 cells per perturbation) rather than a summary of the response (pseudobulk expression, list of DE genes, etc.) for two reasons. First, single cells yield every Perturb-seq result (distances, differential expression, pathway enrichment, gene signatures, etc.), while collecting each result separately would invite a different model for each outcome, which is less likely to generalize. Second, if we want a model to stand in faithfully for the experiment, the model has to predict what the experiment measures, including its noise and biases.

We also require raw counts rather than log-normalized values. In the 2025 Challenge, the scorer accepted either, without any limit on their scale, and shortcut solutions exploited this freedom to rescale the data to unrealistic values. Integer counts, with at most 10^6^ per cell, fix the space and bound the scale. Finally, the predictions are compared with the measured control cells of the reference, rather than with control cells predicted by the submission. In the 2025 Challenge, the metrics used the control cells of the submission and shortcut solutions exploited this freedom as well.

## 3. Shared design of the six metrics

### 3.1 Target genes: excluded from every metric

Every knockdown lowers the expression of its own target gene, and the participants were given the list of targets. Predicting this drop requires no model, so we exclude the target gene of each perturbation from all six metrics. The perturbation-discrimination metric compares the prediction for each perturbation with the measured profiles of all perturbations, which show the knockdowns of their own targets. For this metric, we remove all 300 target genes of the panel from every profile, so that all comparisons use the same genes. In the 2025 Challenge, the target gene was excluded only from this metric.

### 3.2 Scores: anchored at the mean baseline and at a repeat of the experiment

The raw values of the six metrics cannot be compared with each other. They differ in range (Fig. 3.1), and the values that can be reached differ between the cell lines. To place the metrics on one scale, we anchor each of them at two values measured on the same reference data: a mean baseline *b*, the value of a prediction without any perturbation-specific information, and a replicate *r*, the value of a repeat of the experiment. Like the measured responses to the perturbations, the per-line values of the two anchors were not released to the participants. A submission’s value *u* of a metric becomes the score

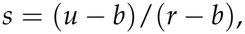

so that 0 is the mean baseline and 1 is a repeat of the experiment. The score of a line is the equal-weight mean of its six scores, and the scores of the lines are averaged. We weight the six scores equally, since the anchoring places them on one scale and each of them scores a different property of the prediction. Sections 4 and 5 give the values of the metrics before this scaling.

**Fig. 3.1.**
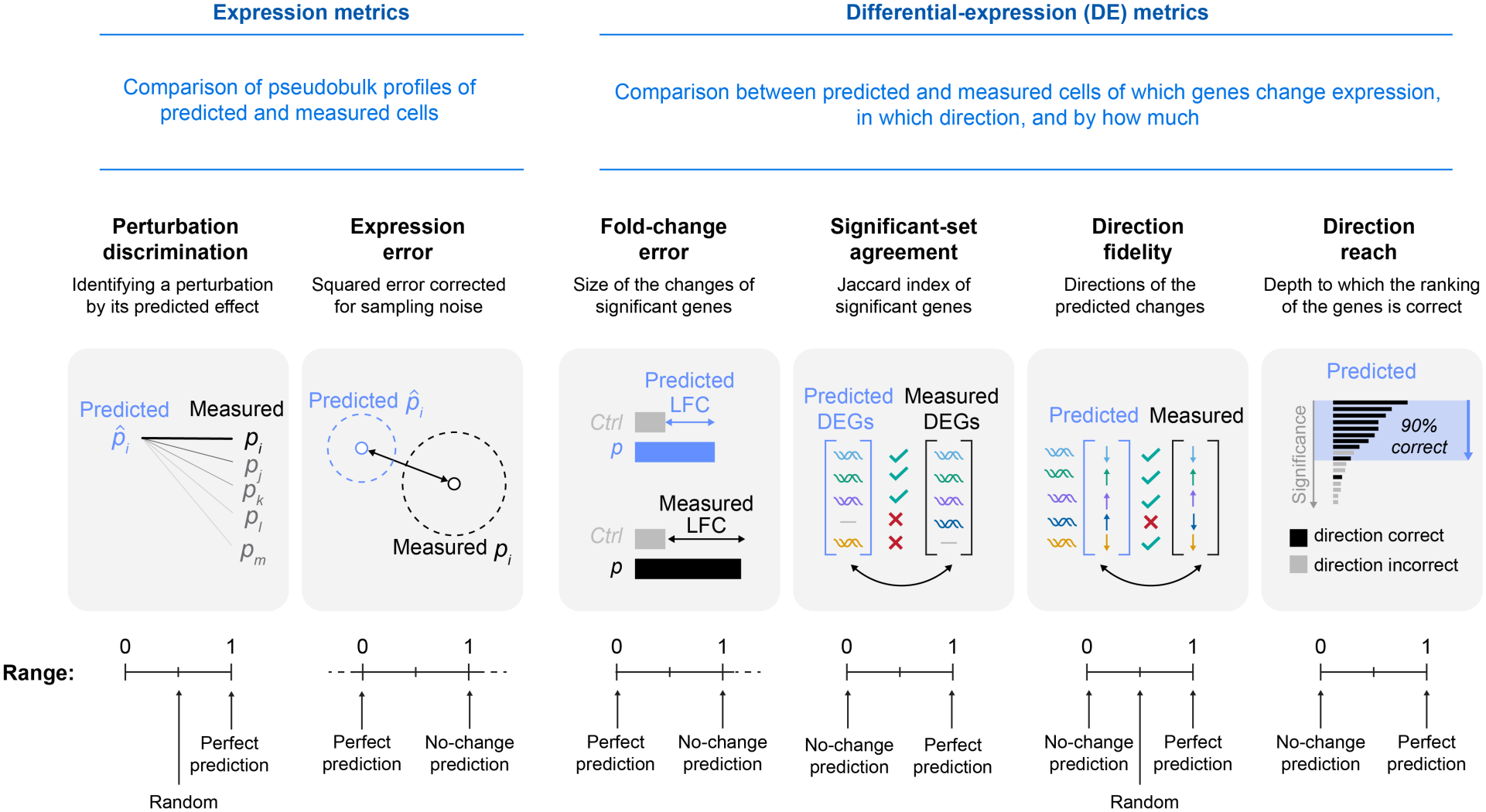
The six metrics of the 2026 Challenge.

The mean baseline is the mean response of the line: the equal-weight average of the mean count profiles of all constructs screened in the line with at least 300 cells that pass a knockdown filter (4,497–6,010 constructs per line, most of them outside the panel). It is assigned to every perturbation as 400 identical cells and scored as an ordinary submission, except that its counts are fractional. The replicate compares the reference with itself. We split the cells of every perturbation and of the control into two disjoint halves, scored one half against the other, each with its own control cells, and averaged the result over five random splits. For the fold-change error, only the fold changes came from the halves, while the significant genes and the denominator came from the full reference. We note that Schäfer et al. (2026) discuss the same normalization, between a lower reference and a positive reference made by splitting the cells of a perturbation into two disjoint subsets (see also Vollenweider and Bühlmann, 2026). As Schäfer et al. point out, such a normalization is interpretable only if the replicate scores better than the mean baseline, which holds for all six metrics on all six cell lines. Since every half holds only 200 cells per perturbation, the replicate is noisier than the full reference, and a prediction equal to the reference scores above 1. A model that predicts the response exactly can therefore also score above 1, by a margin that differs between the metrics. We do not clamp the scores above 1 or the negative scores of submissions worse than the mean baseline. The two exceptions are the two error metrics, which have no upper bound, so that one badly scaled prediction could otherwise dominate the mean score. The score of the fold-change error is floored at −6. The score of the expression error is clamped to the range from 0 to 1, since its corrected error can also fall below that of the replicate, and even below zero, without a better prediction.

In the 2025 Challenge, the upper anchor was the ideal value of each metric (zero for an error metric, one for the others), and scores below the mean baseline were set to zero. We replaced the ideal by the replicate because no prediction can be expected to reach the ideal value: even a repeat of the experiment falls short of it, owing to the measurement noise. At the lower end, clamping at zero hid how far below the mean baseline a submission fell, which is why negative scores are now included for five of the six metrics.

## 4. Expression metrics

Both expression metrics (perturbation discrimination and expression error) compare the pseudobulk profiles of predicted and measured cells. In the 2025 Challenge, the pseudobulk profile was calculated as the mean over the cells of their log-normalized expression. Since the logarithm is concave, this mean (log-norm) depends on how the counts are spread over the cells (Jensen’s inequality), and a prediction with the correct expression but with less variation between its cells than the measured cells cannot reach the measured profile, however accurate it is. In the 2026 Challenge, we form the profile of a group of cells (the cells of one perturbation, or the control cells) by summing the counts of each gene over the cells of the group, normalizing the sums to a fixed total TS = 5 × 10^4^, and log-transforming them:

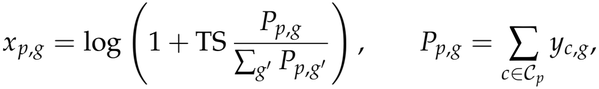

where C_*p*_ is the set of cells of perturbation *p*, and *y*_*c,g*_ is the count of gene *g* in cell *c*.

The effect of a perturbation *p* is its profile minus the profile of the measured control cells, *δ*_*p*_ = *x*_*p*_ − *x*_ctrl_. We mark predicted profiles and effects by a hat, and form the predicted effects against the same measured control. Both metrics use all 18,533 genes except the target genes, and do not filter out low-expressed genes.

### 4.1 Perturbation discrimination: identifying a perturbation by its predicted effect

The perturbation-discrimination metric asks whether a prediction identifies its perturbation: is the predicted effect of a perturbation closer to the measured effect of the same perturbation than to the measured effects of the other perturbations? It is the transposed retrieval rank of Wu et al. (2024), as Schäfer et al. (2026) call it, and it was one of the three metrics of the 2025 Challenge (Roohani et al., 2025).

We compare the predicted effect of each perturbation *p* with the measured effects of all *n* = 300 perturbations of the line by the cosine distance

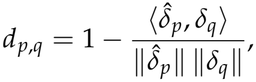

computed without the 300 target genes of the panel, so that all *n*^2^ distances use the same genes. If either vector has zero norm, *d*_*p,q*_ = 1. We score perturbation *p* by its perturbation discrimination score (PDS), one minus the normalized rank of its own measured effect among the *n* candidates,

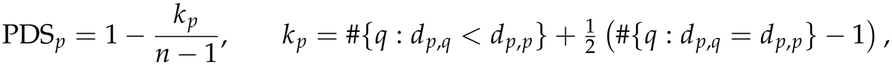

where tied candidates share the mean of the ranks they span, and we average PDS_*p*_ over the 300 perturbations of a line. The score ranges from 0 to 1, and a prediction without information on the identity of the perturbations scores 0.5 on average.

We revised the perturbation-discrimination metric from the 2025 Challenge in two ways. First, we replaced the L1 distance by the cosine distance. The L1 distance depends on the length of the predicted effect as well as on its direction, so that the score of a prediction changes when its effects are rescaled (Supplementary Note 2 and Supplementary Fig. S2), as Liu et al. (2025) showed. Second, we removed all 300 target genes of the panel rather than only the target of the predicted perturbation. If only that target is removed, the measured effect of every other perturbation still shows the knockdown of its own target. A prediction that raises the expression of the other 299 target genes then ranks its own perturbation first, using only the list of target genes.

On the six cell lines, the half-replicate scores 0.923–0.984, and it falls short of 1 almost exclusively on perturbations with weak effects (Fig. 4.1). On every line, 99–100% of its shortfall comes from perturbations whose effect is smaller than their sampling noise, and perturbations with an effect below a tenth of their noise score 0.74 on average.

**Fig. 4.1.**
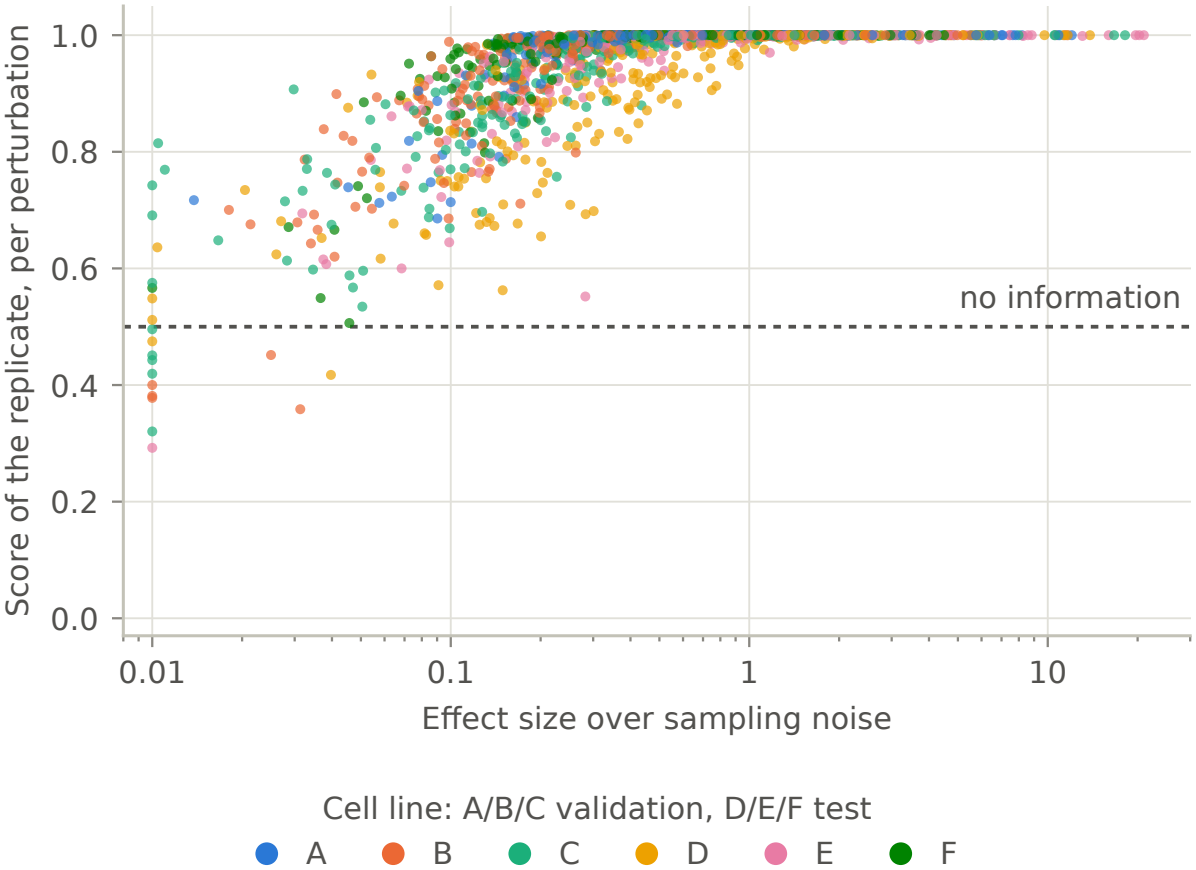
A repeat of the experiment fails to identify only the perturbations with weak effects. PDS of each perturbation for the replicate (half of the cells of every perturbation predicting the other half, each half with its own control cells), mean of the five splits, against the effect size of the perturbation over its sampling noise: the squared distance between the profiles of the perturbation and of the control, corrected for sampling noise, divided by the noise of the two profiles, from all cells of the line. Effect sizes below 0.01 (16 of 1,800 perturbations, 15 of them at or below 0) are drawn at 0.01.

### 4.2 Expression error: the squared error corrected for sampling noise

The expression error measures how far the predicted profile is from the measured one. The plain mean squared error between the two profiles does not measure this distance faithfully. Both profiles are estimated from finite samples of cells, and the sampling noise of each inflates the squared distance between them. For two independent estimates of the profiles, the expected squared distance between them is the squared distance between their expected values plus the sampling variances of both, summed over the genes:

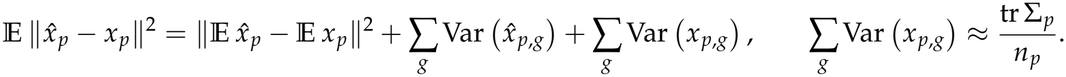

The second relation holds exactly for a profile that is the mean over *n*_*p*_ independent cells whose expression has the covariance matrix Σ_*p*_, and approximately for our profiles. It shows that the inflation decreases only in inverse proportion to the number of cells. Subtracting unbiased estimates of the two variance terms gives an unbiased estimate of the squared distance between the expected profiles. This correction underlies the high-dimensional two-sample tests of Bai and Saranadasa (1996) and Chen and Qin (2010), whose statistics are unbiased estimates of the squared distance between two means.

At the depth of the Challenge data, the sampling noise makes up 38–60% of the squared distance between the measured profiles of the perturbations and of the control. For the individual perturbations, a repeat of the experiment, in which half of the cells of every perturbation predict the other half, has a larger uncorrected squared error than a no-change prediction (i.e. pasting the profile of all 18,400 control cells as the prediction) for 78–94% of the perturbations (Fig. 4.2a).

**Fig. 4.2.**
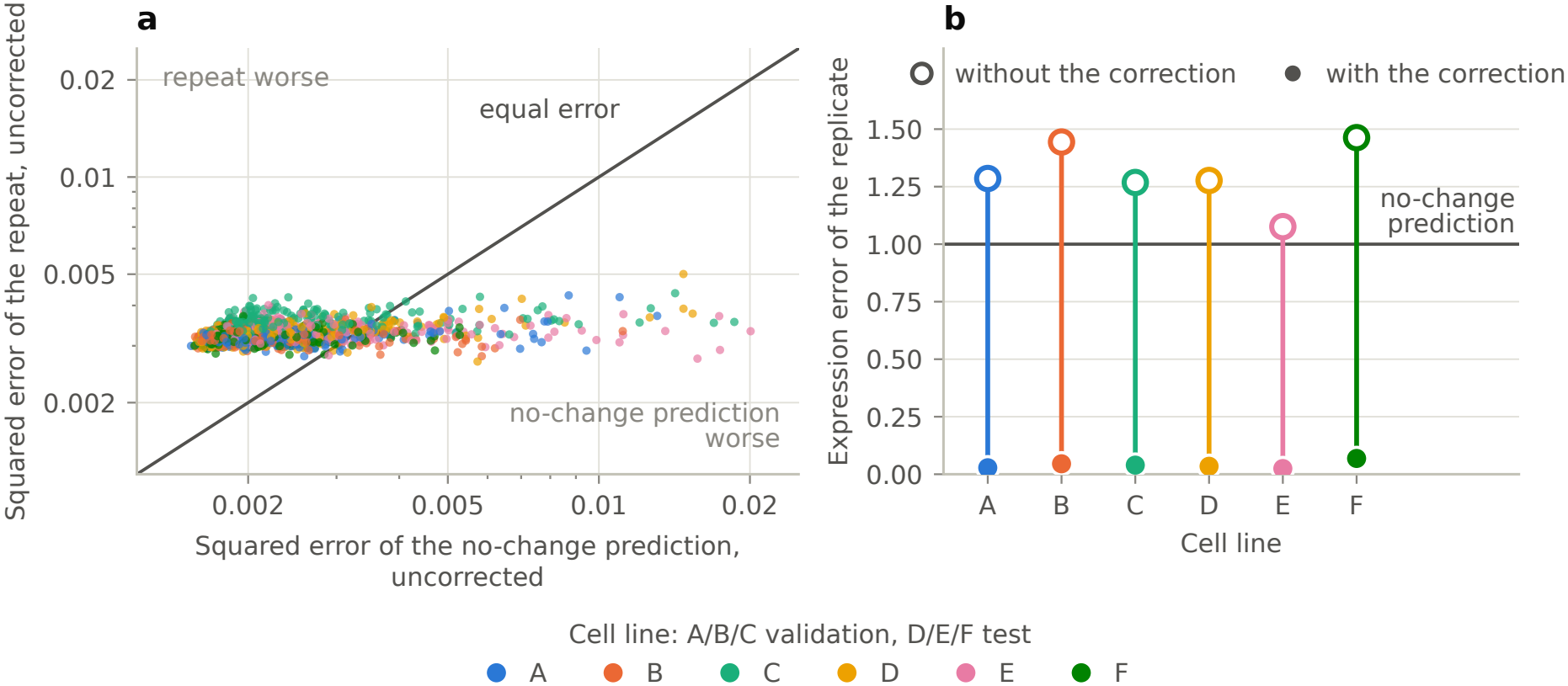
Without its sampling correction, the expression error ranks a no-change prediction above a repeat of the experiment. a: Squared error of each perturbation without the correction, for a repeat of the experiment (half of the cells of every perturbation predicting the other half) and for a no-change prediction (the profile of all 18,400 control cells), both against the same measured half, mean of the five splits, log scale. Points above the diagonal are perturbations for which the repeat scores worse than the no-change prediction. **b:** Expression error of the replicate (the same repeat, each half with its own control cells), mean of the five splits, per cell line, without (open circles) and with (filled circles) the sampling correction.

Over all perturbations of a line, the uncorrected error of the replicate exceeds the error of the no-change prediction on all six lines (Fig. 4.2b, 1.08–1.46 versus 1). Without the correction, a repeat of the experiment would thus score worse than the no-change prediction. Schäfer et al. (2026) pointed out this problem: on profiles over all genes, the uncorrected squared error often fails to distinguish a positive reference from a negative reference, in their case the mean of all perturbed cells. With the correction, the replicate scores 0.025–0.069, and the no-change prediction 1.01–1.03.

The mean absolute error of the 2025 Challenge, over all genes and without any correction, suffered from the same problem. On our six lines, with the pseudobulk profiles of the 2025 Challenge, the absolute error of a repeat of the experiment exceeds that of the no-change prediction as well, by a factor of 1.10–1.25 on every line.

We estimate the sampling noise of a profile by the delete-one jackknife approximation (Efron and Stein, 1981):

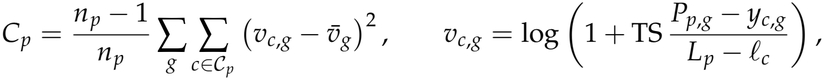

where *n*_*p*_ is the number of cells of the group, *L*_*p*_ = ∑_*g*_ *P*_*p,g*_ is its total count, *l*_*c*_ is the total count of cell *c*, and the bar denotes the mean over the cells of the group. *C*_*p*_ estimates the part of a squared distance that remains even when the two compared profiles agree in expectation. We compute it in the same way for the prediction, *Ĉ*_*p*_, and for the control cells, *C*_ctrl_.

As the numerator of the error, we take the corrected squared distance between the predicted and measured profiles,

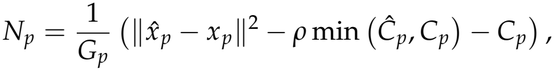

where *G*_*p*_ is the number of genes summed (all genes except the target gene of *p*). The minimum and the factor *ρ* ≤ 1 are introduced to prevent a shortcut solution from claiming more credit by inflating its noise. The minimum caps the correction credited to the prediction at the correction of the measured profile, so that emitting cells more variable than the measured cells does not lower the error. The factor *ρ* caps the total correction of a submission by the variation of its predicted profiles across the perturbations:

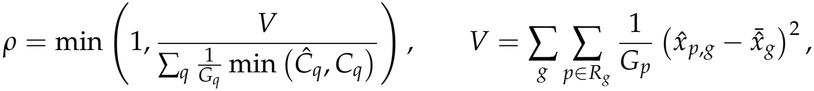

where *R*_*g*_ is the set of perturbations that score gene *g* (all perturbations except the one targeting *g*), and the bar denotes the mean over *R*_*g*_, weighted by 1/*G*_*p*_.

The denominator of the expression error metric is the corrected squared distance between the measured profile and the control profile,

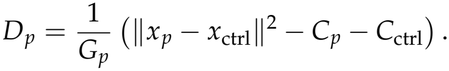

Dividing by it expresses the error as a fraction of how far the measured profile is from the profile of the control cells. This follows the common practice of reporting a squared error relative to that of a no-change prediction. We sum both distances over all genes except the target gene of the perturbation. We subtract the corrections whole, including the small share of the excluded gene. This changes the sum of the denominators by less than 0.02%. The error of a line is the ratio of the two sums over its perturbations,

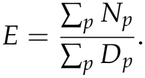

We take the ratio of the sums rather than the mean of per-perturbation ratios, because the corrected denominator of a perturbation with a weak effect is close to 0 and can be negative, which makes its ratio unstable. The denominator depends on the measured data alone, and is the same for every submission scored on a line. When all denominators are positive, the error of a line is the weighted mean of the per-perturbation ratios, with their denominators as weights, so that the perturbations with strong effects weigh more.

In summary, our expression error is based on a nearly unbiased estimate of the squared error, which corrects for the sampling noise that otherwise dominates the squared error of an accurate prediction. Two additional measures, the minimum and the factor *ρ*, are designed to prevent shortcut solutions from taking extra credit for the noise.

Despite our efforts to prevent shortcut solutions, we believe that they are still possible, and we have evidence that they occurred in the 2026 Challenge. Both min(*Ĉ*_*p*_, *C*_*p*_) and *ρ* cap the noise credit, but neither can verify that a predicted profile carries the noise for which it is credited. On the final validation leaderboard, all ten leading submissions collected 91–100% of the full credit, the largest correction that the minimum allows. This credit is fair only when the predicted profiles carry that much noise, as they do when the submitted cells are sampled. Eight of the ten, however, emitted cells whose summed counts, for many genes, varied less across perturbations than sampling allows. With the sums of every perturbation scaled to the median total of the perturbations, their variance divided by their mean, the Fano factor, was below 0.5 for 26–50% of a line’s genes with a mean of at least 10 counts, whereas Poisson-sampled sums with the same gene means keep it above 0.6 for every gene. Their predicted profiles therefore carried less noise than they were credited for. Averaged over genes, at least 23–41% of the Poisson sampling noise was missing from their summed counts.

## 5. Differential-expression metrics

The main readout of a perturbation experiment is the list of genes that respond to each perturbation: which genes change their expression, in which direction and by how much. These differentially expressed genes are the biologically relevant information that we want to extract from the experiment, since they point to the function of the perturbed gene and to the pathways and programs that it controls. A model that predicts the outcome of the experiment therefore has to reproduce them, and four of the six metrics compare the DE genes of a submission with those of the measured data. In the 2025 Challenge, a single metric scored the DE genes: the overlap between the measured significant genes and the predicted significant genes with the largest absolute fold changes (at most as many as the measured ones), relative to the number of measured ones (Roohani et al., 2025). We replaced it with four metrics, which score the size of the changes, the set of significant genes and the directions of the changes.

To find the DE genes, we compare the cells of each perturbation with the measured control cells, gene by gene, by the two-sided Wilcoxon rank-sum test. The test makes no assumption about the distribution of the counts, which are sparse and overdispersed, and an outlying cell affects it only through its rank, not through its value. We apply the test to the counts normalized per cell, and take its p-values from the normal approximation with tie and continuity corrections, which is accurate at the sizes of the groups (400 cells per perturbation, 18,400 control cells). We correct the p-values for multiple testing within each perturbation by the Benjamini–Hochberg procedure (Benjamini and Hochberg, 1995) and call a gene significant if its adjusted p-value is below 0.05, i.e. at a false discovery rate of 5%. The test reports only whether a gene changed. The direction and the size of the change come from the log_2_ fold change,

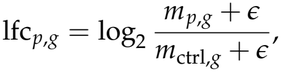

where *m*_*p,g*_ and *m*_ctrl,*g*_ are the means of the normalized counts (to a total of 10^6^ per cell) of gene *g* over the cells of perturbation *p* and over the control cells, and *ϵ* = 10^−9^ keeps the ratio finite when a mean is 0. We apply the same procedure, with the same parameters and against the same measured control cells, to the submission and to the reference. Compared with the 2025 Challenge, we correct for multiple testing within each perturbation rather than across all perturbations together, so that the threshold has the same meaning for a perturbation with a strong response as for one with a weak response. We also compute the fold changes from arithmetic rather than geometric means.

For the DE metrics, we filter out the genes with low expression, testing a gene only if its mean expression in the control cells exceeds 5 counts (UMIs) per million (CPM). This filter keeps 9,537–10,124 of the 18,533 genes, depending on the cell line. To avoid shortcut solutions, we apply the filter to the control cells alone. The tested genes are then the same for every perturbation and for the submission and the reference, and their selection reveals nothing about the response. In this way, we miss the pairs of a perturbation and a gene in which the gene is expressed below the cutoff in the control cells but above it in the cells of the perturbation. Such pairs make up ~1% of all pairs that pass the cutoff in either group.

### 5.1 Fold-change error: the size of the changes

The fold-change error measures whether a submission predicts the size of the changes of the significant genes, which was not captured by any metric in the 2025 Challenge. For each perturbation *p*, we take the set *S*_*p*_ of the genes that the reference calls significant and for which it gives a finite fold change, and compute the mean absolute error of the predicted fold changes over these genes, normalized by the mean absolute measured fold change:

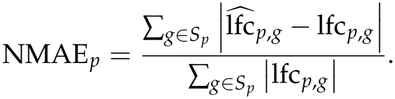

We score only the perturbations with at least 10 genes in *S*_*p*_, since an error over a handful of genes close to the threshold is mostly noise. The error of a line is the mean of NMAE_*p*_ over these perturbations.

A perfect prediction scores 0, and a no-change prediction scores exactly 1, since its error equals the denominator gene by gene. The mean baseline scores 0.99–1.12, i.e. its fold changes are not appreciably closer to the measured ones than those of the no-change prediction. The replicate scores 0.36–0.45 (Fig. 5.1a): half of the cells reproduce the fold changes of the other half with ~40% of the error of the no-change prediction.

### 5.2 Significant-set agreement: the Jaccard index of the significant genes

The significant-set agreement asks whether a submission calls the same genes significant as the measurement. It is the Jaccard index of the two sets of significant genes,

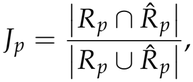

where *R*_*p*_ is the set of genes that the reference calls significant for perturbation *p*, and the set marked by a hat is that of the submission. Unlike the overlap of the 2025 Challenge, which was a fraction of the measured significant genes, the Jaccard index charges every falsely called gene as much as a missed one. If neither the submission nor the reference calls any gene, *J*_*p*_ = 1. The agreement of a line is the mean of *J*_*p*_ over its 300 perturbations.

The Jaccard index of a repeat of the experiment is low. The replicate scores 0.35–0.43 (Fig. 5.1b), i.e. on average the significant sets of the two halves of the data share fewer than half of the genes in their union. The significant set is thus poorly reproducible, even when the experiment itself is reproducible. This is a property of significance testing and of its limited power at these numbers of cells, rather than a failure of the experiment or of the metric. A gene is called significant when its test statistic crosses a threshold, and a gene whose expression truly changes crosses it with a probability (the power of the test) that is far below 1 when the change is small compared with the sampling noise. Two repeats of the experiment therefore call different subsets of the responding genes.

**Fig. 5.1.**
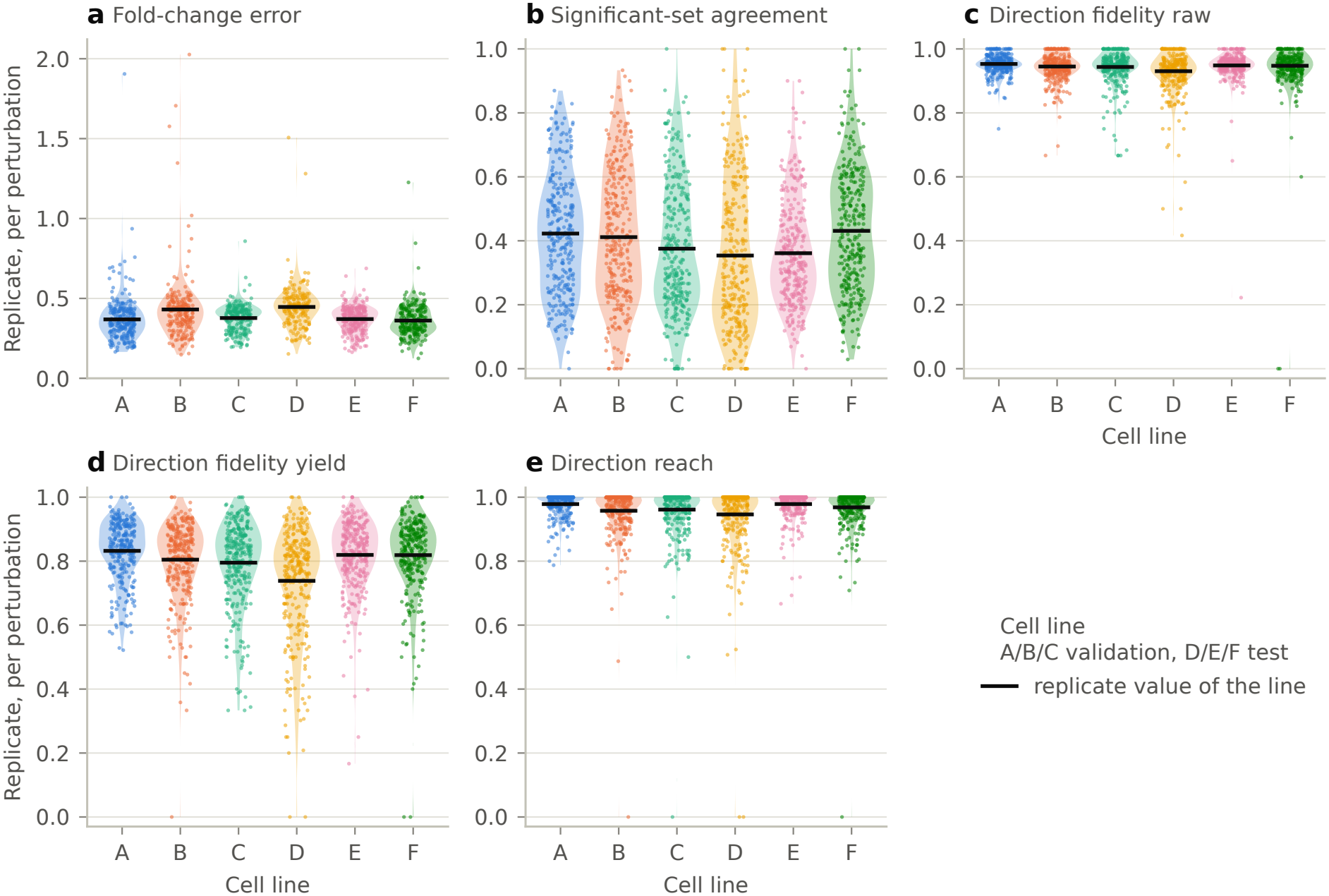
The DE metrics of a repeat of the experiment, per perturbation. Each point is a perturbation, scored for the replicate (half of the cells of every perturbation predicting the other half, each half with its own control cells), mean of the five splits in which the metric is defined. The violins show the distribution over the perturbations of a cell line, and the black bars the value of the replicate for the line. **a:** Fold-change error over the 188–262 perturbations per line that it scores. **b:** Significant-set agreement. **c:** Direction fidelity raw, which is not scored. **d:** Direction fidelity yield. **e:** Direction reach.

### 5.3 Direction fidelity: the directions of the predicted changes

To depend less on the poorly reproducible significant sets, we introduce the direction fidelity metric. Instead of asking whether a submission calls the same set of genes as the reference, it asks whether the predicted direction of gene change (up or down) agrees with the measured direction. A gene that the submission calls significant is judged by the sign of its measured fold change, whether or not the reference calls it significant. For perturbation *p*, let *n*_pred_ be the number of genes that the submission calls significant and whose measured fold change is defined and not 0, and *k* the number of them whose predicted and measured fold changes have the same sign. The direction fidelity raw is the fraction of correct directions,

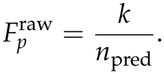

The direction fidelity raw of the replicate is high, 93–95%, close to the 97.5% expected when 5% of the calls are false (Fig. 5.1c). The direction fidelity raw alone invites a shortcut solution: a submission could call significant only the few genes whose direction it is most sure of, and reach a direction fidelity raw close to 1 while saying nothing about most of the response. To prevent it, we score the direction fidelity yield, which divides *k* by the larger of *n*_pred_ and the number *n*_real_ of genes that the reference calls significant:

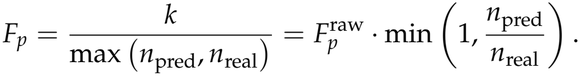

The second form shows that the score is the direction fidelity raw multiplied by a coverage term, which is 1 when the submission calls at least as many genes as the reference and falls in proportion when it calls fewer. The significant set of the reference enters only through its size, not through the genes that it holds. A submission that calls nothing scores 0 whenever the reference calls any gene, and one that calls every gene significant with random directions scores 0.5. The direction fidelity yield of a cell line is the mean of *F*_*p*_ over its perturbations.

The mean baseline, which calls most of the tested genes significant with the same directions for every perturbation, scores 0.45–0.52, at the chance level of 0.5. The replicate scores 0.74–0.83 (Fig. 5.1d).

We appreciate that the current definition of the direction fidelity yield penalizes under-prediction more than over-prediction. A submission that calls fewer genes than the reference loses in proportion, even if all its directions are right, while one that calls more genes loses only through the wrong directions among its additional calls. We accept this asymmetry, because the penalty for calling too few genes is what closes the shortcut solution above.

### 5.4 Direction reach: the depth to which the ranking of the genes is correct

Whether the submission calls a gene significant depends on the calibration of its adjusted p-values, i.e. on how much its cells vary within a perturbation. Correct calibration is a hard task, since it requires accurate modeling of the experimental noise. The direction reach requires no calibration of the false discovery rate, only a correct ranking of the genes by their significance. We order the genes that the reference calls significant for perturbation *p* by the submission’s own significance (by its adjusted p-value, then by its unadjusted p-value, then by its absolute fold change), and find the deepest prefix of this ranking in which at least 90% of the predicted directions are correct:

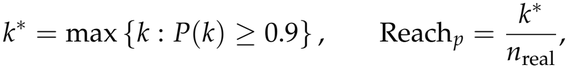

where the purity *P*(*k*) is the fraction of correct directions among the first *k* genes whose measured fold change is defined and not 0, and *k*^*^ = 0 if no prefix reaches 0.9. The purity as a function of the depth *k* of the ranking is the purity curve, and *k*^*^ is the deepest point at which this curve is at or above 0.9. The reach is thus the fraction of the measured DE genes that the submission ranks, from its most significant call down, with at most one wrong direction in ten. A submission that does not calibrate its false discovery rate but ranks the genes correctly can score low on the direction fidelity yield and still high on the reach. The reach of a line is the mean of Reach_*p*_ over the perturbations for which the reference calls any gene significant. The mean baseline scores 0.05–0.10, and the replicate 0.95–0.98, close to the maximum of 1 (Fig. 5.1e).

## 6. Discussion

In this work we present the six metrics that score the 2026 Challenge, together with the reasoning behind each of them and their values on the data of the Challenge. Two of the metrics compare the pseudobulk expression profiles of the predicted and measured cells, while four compare the genes that respond to each perturbation. Every metric is reported on a scale from a mean baseline to a repeat of the experiment, and every definition includes safeguards against shortcut solutions.

The first lesson of this work is that the details of a metric decide which predictions win. With the L1 distance of the 2025 Challenge, the perturbation-discrimination metric rewarded or penalized a prediction for the scale of its effects (Supplementary Note 2). Without a correction for the sampling noise, both the squared error and the absolute error score a repeat of the experiment worse than a no-change prediction (Section 4.2). Such properties are easy to overlook in the definition of a metric. They become obvious when the metric is applied to predictions whose quality is known in advance. Metrics therefore have to be designed as carefully as the experiments whose predictions they evaluate.

The most informative of these predictions is a repeat of the experiment, which we made by splitting the cells of every perturbation and of the control into two halves. A repeat predicts the measurement as well as the experiment itself can. Thus, a metric that ranks it below a trivial prediction does not measure what it is meant to measure. The values of the repeat also show how much of the range of a metric the measurement noise leaves within reach. Two halves of the same experiment share fewer than half of the genes in the union of their significant sets (Section 5.2), while 93–95% of the significant calls of one half have the direction measured in the other (Section 5.3). We therefore anchored every score at the repeat (Section 3.2), and we recommend evaluating any new metric on half-splits of the data before it is used to rank models.

Another lesson is that the metrics should follow the experiment. Four of the six metrics compare the DE genes, the main readout of a Perturb-seq experiment: which genes change, in which direction and by how much (Section 5). A prediction that scores well on these metrics reproduces the result that an experimentalist would report, rather than a summary statistic that is convenient to compute. We believe that the evaluation of virtual cells should be built on the quantities that biologists measure in their experiments. The same principle guides the choice of metrics in biomedical image analysis (Maier-Hein et al., 2024).

Finally, the metrics of a competition have to be designed against shortcut solutions, since a competition rewards whatever raises the score. The tested genes are chosen from the control cells alone, and the target genes are excluded. The predictions are submitted as counts of single cells, and compared with the measured control cells. The expression error caps the credit for sampling noise, and the direction fidelity yield penalizes a submission that calls too few genes.

The new set of metrics improves upon those used in the 2025 Challenge in three ways. First, the DE genes are scored by the size and the direction of their changes, and not only by the overlap of the significant sets (Section 5). Second, a repeat of the experiment scores above the mean baseline on all six metrics and all six cell lines (Section 3.2), while the mean absolute error of the 2025 Challenge scored it below a no-change prediction (Section 4.2). Third, the shortcut solutions that the metrics of the 2025 Challenge left open can no longer raise the score (Sections 2.3, 3.1 and 4.1).

However, there is still room for improvement to prevent shortcut solutions. On the final validation leaderboard of the 2026 Challenge, eight of the ten leading submissions emitted cells whose summed counts, for many genes, varied less than sampling allows. Nevertheless, the expression error credited them with 91–100% of the full correction for sampling noise (Section 4.2). The direction fidelity yield penalizes under-prediction more than over-prediction (Section 5.3), and the significant-set agreement of a repeat of the experiment is low, because the power of the test is limited (Section 5.2). In addition, all six metrics compare the prediction with the measurement within each cell line, and they are not designed to measure how the response to a perturbation depends on the cell line. We find that the leading submissions reproduced only a small part of the measured differences between cell lines (context specificity). After the mean response of each line is removed and the sampling noise is corrected for, about half of the measured response to a perturbation differs between the three validation lines, while the same ten leading submissions reproduced only 2–12% of this line-specific part.

There are multiple areas where our metrics can be improved. We will continue this work for the next Challenge in two directions: metrics that capture more of the biology that experimentalists measure, and designs that close the shortcut solutions that remain open. We believe that carefully designed metrics, calibrated on repeats of the experiment and hardened against shortcut solutions, are as important for building the virtual cell as the models themselves.

## Acknowledgements

This study was supported by the Arc Institute and NVIDIA Corporation.

## Use of generative AI

Claude Code (Anthropic) and Codex (OpenAI) were used to assist in analyses and manuscript writing. The ideas, design decisions, analysis questions, interpretations, final verification and acceptance are the Authors’, who take full responsibility for the content.

## Arc Virtual Cell Team

Usha Adhikari, Will Agnew-Svoboda, Valeria Aizen, Dana AL-Azzeh, Daniel Alameda Serrano, William E. Allen, Alexander Atamian, Jerika J. Barron, Harry Bevir, Steven Boggess, Paul Bunk, Maria Erendira Calleja Cervantes, Cynthia Chen, Yi-Chen Chih, Cyntia Del Valle Gomez, Xiaolong Dong, Miles Duncan, Carmen Gallardo, Luke A. Gilbert, Gabriel Grenot, Emiliano Josue Hergenreder, Jesse Holt, Ben Hsu, Tony J. Hua, Parsa Idehpour, Aarushi Kalaimani, Julia Kazaks, Trinh Lam, David Lara Astiaso, Junzhe Li, Gabriella Martyn, Satoru K. Miura, Harris Nami, Alexander A. Nevue, Charlene Pan, Khushali Patel, Akshaya Rajan, Kheerthivasan Ramachandran, Sneha B. Rao, Lorena Saavedra, Nerea Santana Gonzalez, Kristen Seim, Malvenderjit Jagjit Singh, Tulasi Indrasinh Solanki, Kiran Sriram, Jin Wen Tan, Ipek Tasan, Michael Teng, Ed Tuan, Tushita Wadhawan, Flora Yi, Nick Youngblut, Jamie Zaikov

All members are affiliated with Arc Institute, Palo Alto, CA, USA. Luke A. Gilbert is also affiliated with the Helen Diller Family Comprehensive Cancer Center and the Department of Urology, University of California, San Francisco, San Francisco, CA, USA. William E. Allen is also affiliated with the Department of Developmental Biology, Stanford University School of Medicine, Stanford, CA, USA.

## Supplementary information

### Supplementary Note 1. The cell-eval2 package

We implemented the six metrics of the 2026 Challenge in cell-eval2, an open-source Python package for the evaluation of models that predict the responses of cells to perturbations. The package is a GPU-accelerated reimplementation of cell-eval (https://github.com/ArcInstitute/cell-eval). Four of the six metrics are computed from tests for differential expression, which run on the GPU through the gpudge package (https://github.com/ArcInstitute/gpudge), or on the CPU when no GPU is available. The input files are AnnData files or, for data too large to hold in memory, compressed archives of the cellstream package (https://github.com/ArcInstitute/cellstream), which are scored on the GPU without loading the whole matrix. In July 2026, we compared an early version of cell-eval2 with cell-eval 0.8.1 on 300 perturbations of one Perturb-seq screen. With the conventions of cell-eval 0.6.6 (preset v1), cell-eval2 took 117 s on one NVIDIA H100 GPU to compute the 26 metrics that both packages share, while cell-eval 0.8.1 took 58 min on 16 CPU threads. With its own conventions (preset v2), which test fewer genes, cell-eval2 took 34 s.

The vcc2026 preset of cell-eval2 holds the configurable parameters of the 2026 Challenge metrics, and the metric specification in the repository of the package (docs/vcc2026_metrics) gives the definition of each metric. Both phases of the 2026 Challenge were scored with version 0.16.0. To allow comparisons with the 2025 Challenge, the package also computes its three metrics with the conventions of cell-eval 0.6.6, which scored that Challenge (preset v1, profile vcc).

The package is available from GitHub (https://github.com/ArcInstitute/cell-eval2) and from PyPI (pip install cell-eval2) under the MIT license.

**Supplementary Fig. S1.**
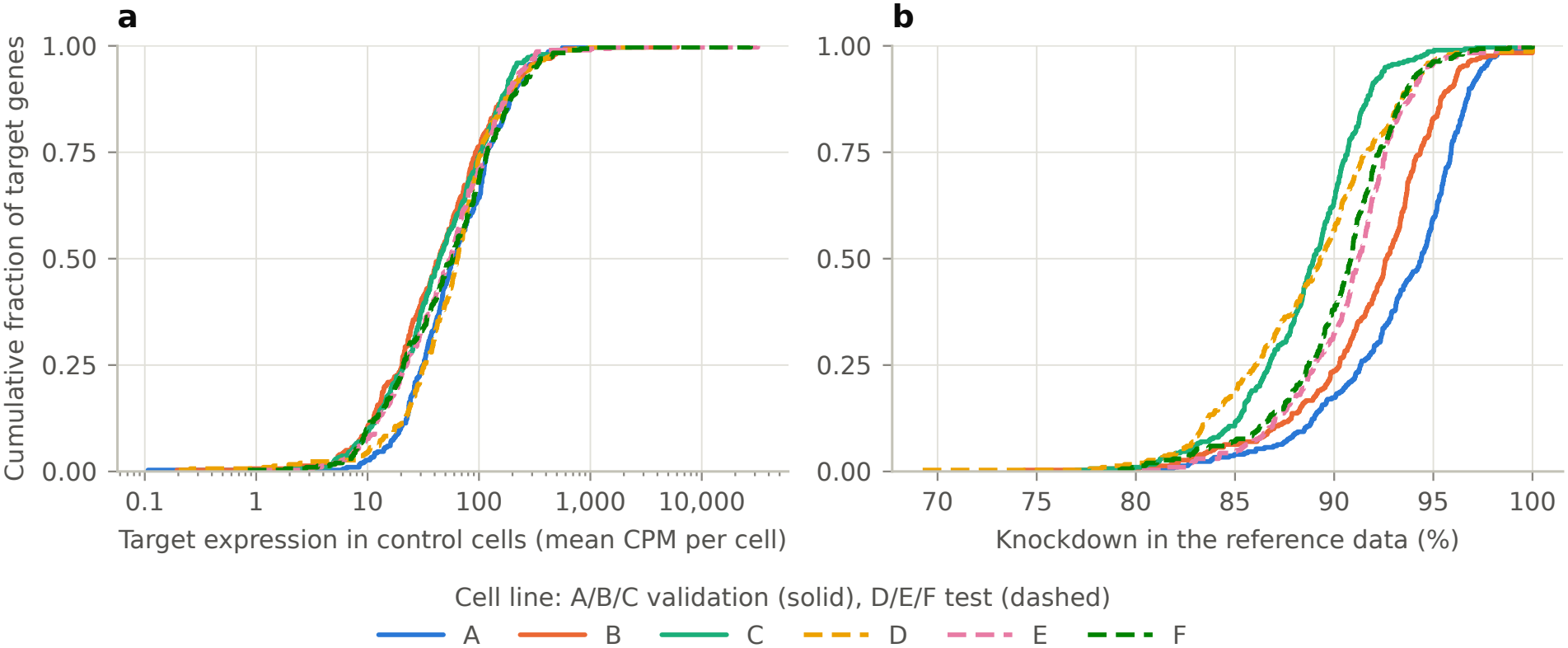
Target expression and knockdown of the panel genes, per cell line (Section 2.2). Cumulative distributions over each line’s 300 target genes. **a:** Expression of the target gene in the line’s 18,400 control cells, the mean over cells of counts per million (CPM), log scale. **b:** Knockdown of the target gene in the reference data, one minus the ratio of its pooled CPM in the perturbation’s 400 cells to its pooled CPM in the control cells, in percent. Pooled CPM is the gene’s summed counts per million summed counts. Solid lines, validation cell lines A/B/C. Dashed lines, test cell lines D/E/F. The three cell lines of a phase share one set of 300 genes.

### Supplementary Note 2. The L1 distance and the scale of the predicted effects

In Supplementary Fig. S2, every predicted effect of the replicate is multiplied by the same factor, and nothing else changes. The directions of the effects stay the same, and with them the information that the prediction carries on the identity of the perturbations. With the cosine distance, which depends on the directions alone, the score is the same at every factor. With the L1 distance of the 2025 Challenge, the score depends on the factor. It falls to 0.79–0.88 when the effects are halved, and to about 0.5, the score of a prediction without information, when they are divided by 64. It is highest when the effects are enlarged by a factor of 1.4–2, slightly above the score of the replicate itself.

**Supplementary Fig. S2.**
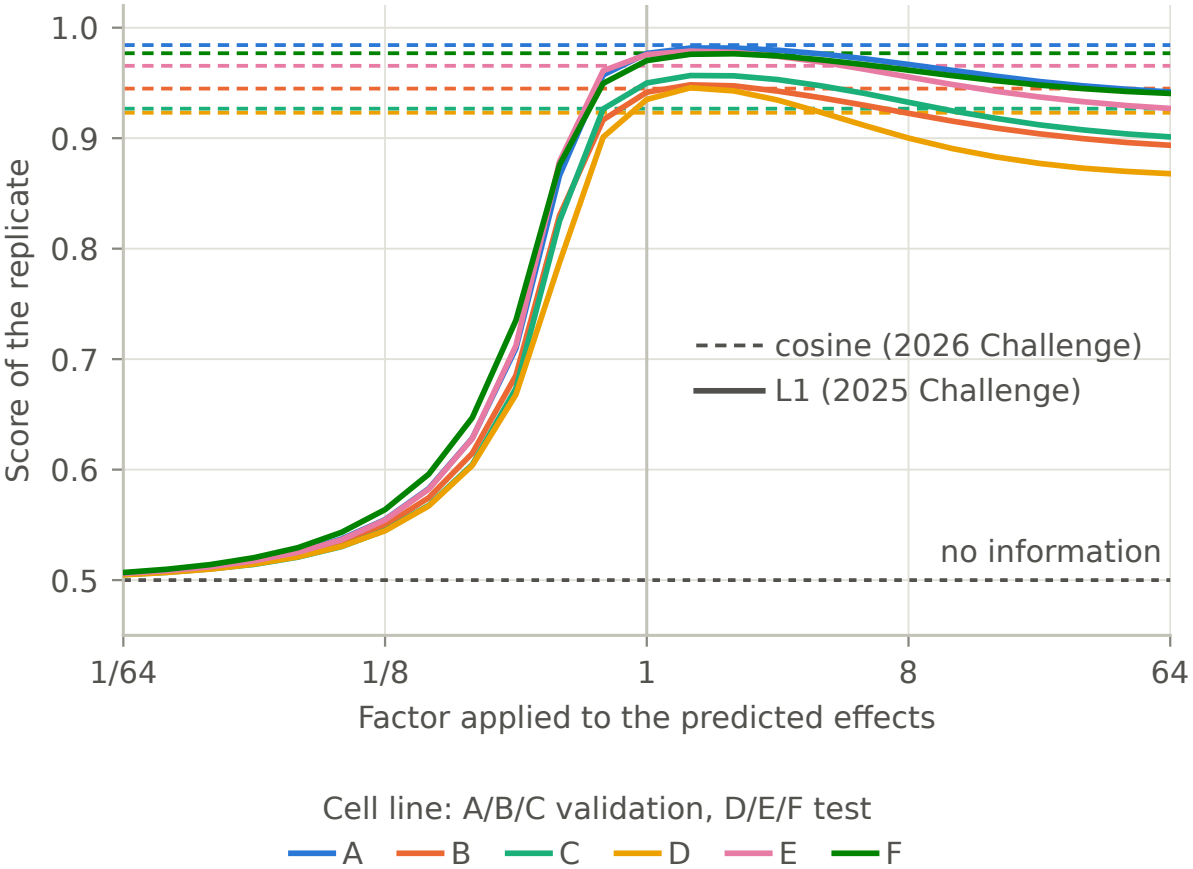
With the L1 distance, the score of the perturbation-discrimination metric depends on the scale of the predicted effects (Section 4.1). Score of the replicate (half of the cells of every perturbation predicting the other half, each half with its own control cells), mean of the five splits, on each cell line when every predicted effect is multiplied by the same factor, with the cosine distance (dashed) and with the L1 distance of the 2025 Challenge (solid), all else as in the 2026 Challenge. A factor of 1 is the replicate itself.

Thus, with the L1 distance, the score of a prediction depends on the size of its effects as well as on how well they identify the perturbations. A prediction with the correct directions is penalized if its effects are too small, and rewarded if they are moderately exaggerated. At the true scale, the two distances score the replicate about equally, 0.923–0.984 with the cosine distance and 0.935–0.977 with the L1 distance. The advantage of the cosine distance is not a higher score for a faithful prediction, but its independence of the scale of the predicted effects, whose size is scored by the expression error instead.

